# Neural correlates of nightmares revisited: findings from large-scale fMRI cohorts

**DOI:** 10.1101/2024.07.02.601684

**Authors:** Mariana Pereira, Noëlle Terpstra, Renate Rutiku, Kristian Sandberg, Martin Dresler, Florian Krause

**Author notes:** Shared senior authorship (Kristian Sandberg, Martin Dresler, Florian Krause). <u>Corresponding author:</u> Mariana Pereira.

## Abstract

**Study Objectives:** Nightmares are linked to daytime distress and psychiatric and neurological disorders. However, knowledge about the brain regions involved in nightmare production is sparse. This study aimed to examine the relationship between nightmare frequency and the functional connectivity of the amygdala with the prefrontal cortex, which is central to emotional regulation. Additionally, the study sought to replicate previous findings on the neural correlates of nightmares using two large cohorts.

**Methods:** 464 participants underwent functional and structural MRI recordings and answered questions assessing nightmare and dream recall frequency. A general linear model assessed the voxelwise correlation between the amygdala-prefrontal cortex connectivity with nightmare frequency. Additionally, regional homogeneity (ReHo) maps were calculated, high and low nightmare frequency and the entire nightmare frequency spectrum contrasts were determined while controlling for age, sex, and dream recall frequency.

**Results:** The study did not find a significant correlation between nightmare frequency and amygdala-prefrontal cortex connectivity. Previous ReHo findings of group differences between high and low nightmare frequency could not be replicated. However, parametric analysis revealed an association between nightmare frequency and ReHo differences in the cerebellum.

**Conclusions:** This failure to confirm hypothesized and previously reported results, especially with a larger sample size, suggests a need to reevaluate the existing knowledge of the neural correlates of nightmares and consider individual differences such as personality, trauma history, and cognitive processes. Overall, our findings highlight the complexities of interpreting neuroimaging data in sleep research.

## Introduction

Nightmares are dreams characterized by strong negative emotions that can cause significant distress and affect daily functioning, with their frequency varying substantially within the population and the precise mechanisms underlying their formation remaining elusive. Nightmares have the potential to disrupt sleep by causing awakenings with vivid recollections of the unpleasant mentation (Nielsen & Levin, 2007; Zadra et al., 2006). Nightmares are common, however their frequency is substantially heterogeneous within the population (American Psychiatric Association, 2013): 85% of adult respondents reported having had a nightmare at least once a year, and about 2-6% reported suffering from weekly nightmares (Levin & Fireman, 2002; Zadra & Donderi, 2000). Some risk factors have been identified in people who experience nightmares frequently, including genetic predisposition (Hublin et al., 1999), state or trait anxiety (Schredl, 2003), increased stress (Picchioni et al., 2002; Schredl & Goeritz, 2019), psychopathologies such as schizophrenia (Levin, 1998), major depressive disorder and bipolar disorder (Akkaoui et al., 2020), and post-traumatic stress disorder (Campbell & Germain, 2016; Harvey et al., 2003; Ross et al., 1989). Despite the clinical relevance of diagnosing and treating nightmares, the exact mechanisms responsible for their formation remain unclear. Consequently, investigating the neurophysiological factors that contribute to nightmare frequency may significantly improve our comprehension of their underlying causes.

Models of nightmare production and emotion regulation emphasize the functional interplay between the amygdala and prefrontal cortex regions, key regions involved in adaptive emotion processing (Berboth & Morawetz, 2021; Nielsen & Levin, 2007). While theoretical debates persist regarding the extent of emotional regulation during sleep and its manifestation in dreams, accumulating evidence suggests that fear experiences in dreams can influence adaptive responses to threats in waking life (Sterpenich et al., 2020). However, the transition from adaptive dream mechanisms to impaired daily functioning, sleep disturbance, and vulnerability to psychological disorders remains unclear. The "Affect Network Dysfunction" model proposes that nightmares result from dysfunction within a brain network that oversees the adaptive function of fear extinction during dreaming (Nielsen & Levin, 2007). Neurobiologically, the basolateral amygdala is critical for fear encoding and extinction, whereas the medial prefrontal cortex mediates expression. In addition, the hippocampus and brainstem modulate contextual cues and suppress conditioned fear expression, respectively. This neural circuit involving the anterior hippocampus, amygdala, and prefrontal cortex is thus thought to influence the occurrence and severity of nightmares (Nielsen & Levin, 2007). For emotion regulation, several prefrontal cortex regions consistently interact with the amygdala during emotional down- regulation via reappraisal (Berboth & Morawetz, 2021; Loos et al., 2020). In support of this, altered gray matter volume in the left inferior frontal gyrus is associated with depression in patients with idiopathic rapid eye movement sleep behavior disorder who have elevated negative emotional dreams (Bourgouin et al., 2019). Conversely, bilateral calcification of the basolateral amygdala correlates with more pleasant dreams, suggesting a role for the amygdala in shaping dream emotion, as patients with basolateral amygdala lesions perceive dreams as less negative (Blake et al., 2019).

Effective emotion regulation plays a key role in managing and responding to evolutionary relevant threats and stress experiences, thereby shaping emotional reactivity and overall well-being. The differential susceptibility framework (Carr & Nielsen, 2017) proposes sensory processing sensitivity as a trait relevant to the study of nightmares, with nightmare-prone individuals exhibiting heightened emotional reactivity to both positive and negative stimuli. Studies of nightmare frequency have shown an inverse relationship between nightmare severity scores and regional cerebral blood flow in the right medial frontal gyrus during negative image viewing (Marquis et al., 2019). This finding was partially replicated using functional near-infrared spectroscopy (fNIRS), suggesting a negative association between dysphoric dream distress and frontal activation during negative image viewing (Carr, 2020). In the context of threat perception and emotion regulation circuits, abnormal resting amygdala-prefrontal cortex connectivity has been associated with repeated childhood stress, contributing to heightened threat perception (Ochsner & Gross, 2005). Disturbances in this connectivity may serve as a prelude to heightened emotional reactivity during dream states, and the abnormal connectivity observed in individuals with psychological disorders suggests that resting-state patterns may provide valuable insight into nightmares and serve as a potential predictor of their occurrence.

Few studies have investigated the neural correlates of nightmares from a whole-brain perspective. As a notable example, (Marquis et al., 2021) investigated the relationship between resting-state functional magnetic resonance imaging (fMRI) and nightmare frequency in a sample of 18 frequent nightmare recallers and 18 controls. They found altered regional homogeneity (ReHo, based on Kendall’s concordance coefficient measures of BOLD time series for each and nearest voxels) in various brain regions, including frontal (medial prefrontal and inferior frontal), parietal, temporal, and occipital regions, as well as some subcortical regions such as the thalamus. Their findings partially support previous research by (Shen et al., 2016), who observed increased ReHo in the left anterior cingulate cortex and right inferior parietal lobule in 15 patients with nightmare disorder. However, when comparing nightmare disorder patients and matched controls, the latter study did not observe ReHo changes in the hippocampus and amygdala. These findings suggest that the severity and frequency of nightmares may be associated with altered neural activity in several brain regions, including those involved in emotion regulation. However, findings are inconsistent and limited by small sample sizes and heterogeneous populations.

To unravel the complex mechanisms underlying nightmare frequency, the present study had two primary goals. First, we aimed to elucidate the interaction between amygdala-prefrontal cortex connectivity and nightmare frequency in a large cohort of healthy volunteers. We hypothesized that robust functional connectivity between these regions may serve as a predictor of attenuated nightmare recurrence. We speculated that the strength of this connectivity promotes a more effective down-regulation process that not only contributes to effective emotional regulation during waking hours, but also positively influences nighttime processes. To strengthen the reliability of our results, we replicated our findings in another large independent cohort of subjects. Second, our research aimed to replicate and extend the existing literature on whole-brain correlates of nightmare frequency by employing ReHo analysis with a significantly larger sample size, while controlling for dream recall frequency. We anticipated that the increased statistical power of a larger cohort would provide more robust insights into the neural correlates of nightmare formation. Consistent with our replication efforts, we anticipated a convergence of findings with previous work. To achieve this, we used voxel-wise correlations to assess amygdala-prefrontal connectivity and generate ReHo maps. We then examined the relationship between these measures and nightmare frequency in two independent analyses. Through these efforts, our study aimed to contribute significantly to understanding nightmares, elucidate the intricate neural mechanisms governing their recurrence, and potentially provide a pathway to more effective diagnostic and therapeutic interventions.

## Materials and Methods

### Study Population

The data were acquired in the context of a large multi-site cohort project as part of the EU COST Action CA18106 “The neural architecture of consciousness” (https://neuralarchcon.org/), and is composed of MRI and behavioural data collected from healthy participants at two data collection sites. For Dataset 1, the study was approved by the regional local ethics committee, *De Videnskabsetiske Komitéer for Region Midtjylland*, Denmark. Participants were recruited through the Center of Functionally Integrative Neuroscience (Aarhus University) participant database and local advertisement. For Dataset 2, the study was approved by *the Research Ethics Committee at the Institute of Psychology* and the *Komisja Bioetyczna* of the Jagiellonian University, Krakow, Poland. Participants were recruited through advertisements on various websites of the Jagiellonian University and Facebook. *Dataset 1:* A total of 306 participants gave informed consent to participate in the study and received financial compensation for their time and contribution. From these 306 participants, fMRI data of 269 participants were available, and among those data, nine participants were excluded: five based on incomplete questionnaires and four based on incomplete fMRI data. Hence, data from a total of 260 participants (152 female, mean age of 24.78 ranging from 18-48 years) was used in this work. *Dataset 2*: A total of 302 participants gave informed consent to participate in the study. They received financial compensation for their time and contribution. From these 302 participants, preprocessed and quality-checked fMRI data of 164 participants were available at the time of the analysis. Hence, data from a total of 164 participants (99 female, mean age of 23.31 ranging from 18-40 years) was used in this work. All participants completed an online questionnaire session from home with a total duration of around 70 minutes, including a seven-point rating scale assessing their dream recall frequency (Schredl & Erlacher, 2004) and an eight-point rating scale measuring their nightmare frequency (Stumbrys et al., 2013). The questionnaires were administered in English for Dataset 1 and in Polish for Dataset 2. Participants were instructed to ensure it was completed in an undisturbed environment. The dream recall scale was recoded into units of mornings per week and the nightmare frequency scale into units per month (Stumbrys et al., 2015).

### Data acquisition

As both sites were part of the same consortium, data collection was standardized for both datasets. Two resting-state fMRI runs (12 and 6 minutes) were recorded alongside quantitative multi-parameter mapping (MPM; (Weiskopf et al., 2013)) and diffusion-weighted imaging in one scanning session lasting approximately one hour. In this study, we restricted our analysis to resting-state fMRI and synthetically generated T1-weighted images (see “Structural data” section below for details). Dataset 1 was acquired at a Siemens Magnetom Prisma-fit 3T MR scanner. For each participant 1500 functional volumes were acquired using a echo planar T2*-weighted sequence sensitive to BOLD contrast with a multiband acceleration factor of 6 (TR/TE = 700/33 ms, flip angle = 53°, field of view = 200 × 200 mm, number of slices = 60, slice thickness = 2.5 mm [no gap], in-plane resolution = 2.5 × 2.5 mm). Dataset 2 was acquired at a Siemens Magnetom Skyra 3T MR scanner, with almost identical parameters, with the only differences being the number of functional volumes (1348) and the TR/TE (801/33 ms).

The MPM protocol was implemented based on the Siemens vendor sequence and was identical for both datasets. Three-dimensional (3D) data acquisition consisted of three multi-echo spoiled gradient echo scans (i.e., fast low angle shot [FLASH] sequences with magnetization transfer saturation (MT), T1, and effective proton density (PD) contrast weighting). Additional reference radio-frequency (RF) scans were acquired. The acquisition protocol had the following parameters: TR of PDw and T1w contrasts: 18 ms; TR of MTw contrast: 37 ms; minimum/maximum TE of PDw, T1w and MTw contrasts: 2.46/14.76 ms; flip angles for MTw, PDw and T1w contrasts: 6°, 4°, 25°, respectively; six equidistant echoes; 1 mm isotropic reconstruction voxel size; Field of view 224 ’ 256 ’ 176 mm; AP phase encoding direction; GRAPPA parallel imaging speedup factor of 2; T1w, PDw and MTw acquisition times: 3:50, 3.50, 7.52. The acquisition of low-resolution 3D spoiled gradient echo volumes was executed using both the RF head coil and the body coil. This dual acquisition facilitated the generation of a relative net RF receive field sensitivity (B1−) map for the head coil 120–122. The approach obtained rapid acquisition by maintaining a low isotropic spatial resolution of 4^3 mm^3^, a short echo time (TE) of approximately 2 ms, and a reduced flip angle of 6°, avoiding parallel imaging acceleration or partial Fourier. This procedure of capturing volume pairs with the head and body coils was systematically repeated before acquiring each of the MT, PD, and T1 contrasts.

### Preprocessing

Data preprocessing was performed using the fMRIprep toolbox version 21.0.2 (Esteban et al., 2019). The toolbox pipeline utilizes a combination of several well-known software packages for fMRI data pre-processing and constitutes a robust tool that also generates quality reports.

#### Structural data

The synthetic T1w images were generated using the longitudinal relaxation rate (R1) and effective proton density (PD) high-resolution maps (acquired during the MPM sequence protocol). First, both maps were thresholded to achieve the required FreeSurfer units. The R1 map was divided by itself two times, thresholded at zero, and multiplied by one thousand. The PD map was thresholded by zero and multiplied by one hundred. All manipulations were performed using *FSL maths* commands. Subsequently, the *mri_synthesize FreeSurfer* command was applied to create a synthetic FLASH image based on the previously calculated T1 (thresholded R1 map) and proton density map. The optional flagged argument for optimal gray and white matter contrast weighting was used with the following parameters 20, 30, and 2.5. Finally, the synthetic T1w image was divided by four according to the scale *FreeSurfer* expected. The pre-processing of the structural data using the *fMRIprep* toolbox was performed in the following steps: firstly, the synthetic T1w images were corrected for intensity non-uniformity (INU) with *N4BiasFieldCorrection* (Tustison et al., 2010), distributed with *ANTs 2.3.3* (Avants et al., 2008, RRID:SCR_004757), and used as T1w-reference throughout the workflow. The T1w-reference was then skull-stripped with a *Nipype* implementation of the *antsBrainExtraction.sh* workflow (from *ANTs*), using OASIS30ANTs as target template. Brain tissue segmentation of cerebrospinal fluid (CSF), white-matter (WM) and gray-matter (GM) was performed on the brain-extracted T1w using *fast* (*FSL* 6.0.5.1:57b01774, RRID:SCR 002823, Zhang, Brady, and Smith 2001). Brain surfaces were reconstructed using *recon-all* (*FreeSurfer* 6.0.1, RRID:SCR 001847 (Dale et al., 1999)), and the brain mask estimated previously was refined with a custom variation of the method to reconcile ANTs-derived and FreeSurfer-derived segmentations of the cortical gray-matter of *Mindboggle* (RRID:SCR_002438 (Klein et al., 2017)). Volume-based spatial normalization to two standard spaces (MNI152NLin2009cAsym, MNI152NLin6Asym) was performed through nonlinear registration with *antsRegistration* (*ANTs* 2.3.3), using brain-extracted versions of both T1w reference and the T1w template. The following templates were selected for spatial normalization: *ICBM 152 Nonlinear Asymmetrical template version 2009c* ((Fonov et al., 2009), RRID:SCR_008796; TemplateFlow ID: MNI152NLin2009cAsym), *FSL’s MNI ICBM 152 non-linear 6th Generation Asymmetric Average Brain Stereotaxic Registration Model* ((Evans et al., 2012), RRID:SCR_002823; TemplateFlow ID: MNI152NLin6Asym0.)

#### Functional data

First, a reference volume and its skull-stripped version were generated by aligning and averaging 1 single-band reference (SBRef). Head-motion parameters with respect to the BOLD reference (transformation matrices, and six corresponding rotation and translation parameters) were estimated before any spatiotemporal filtering using *mcflir*t (*FSL* 6.0.5.1:57b01774, (Jenkinson et al., 2002)). The estimated *fieldmap* was then aligned with rigid-registration to the target EPI (echo-planar imaging) reference run. The field coefficients were mapped on to the reference EPI using the transform. The BOLD reference was then co-registered to the T1w reference using *bbregister* (FreeSurfer) which implements boundary-based registration (Greve & Fischl, 2009). Co-registration was configured with six degrees of freedom. First, a reference volume and its skull-stripped version were generated using a custom methodology of *fMRIPrep*. Several confounding time-series were calculated based on the *preprocessed BOLD*: framewise displacement (FD), DVARS and three region-wise global signals. FD was computed using two formulations following Power (absolute sum of relative motions, (Power et al., 2014)) and Jenkinson (relative root mean square displacement between affines, (Jenkinson et al., 2002)). FD and DVARS are calculated for each functional run, both using their implementations in *Nipype* (following the definitions by (Power et al., 2014)). The three global signals were extracted within the CSF, the WM, and the whole-brain masks. Additionally, a set of physiological regressors were extracted to allow for component-based noise correction (*CompCor*, (Behzadi et al., 2007)). Principal components were estimated after high-pass filtering the *preprocessed BOLD* time-series (using a discrete cosine filter with 128s cut-off) for the two *CompCor* variants: temporal (tCompCor) and anatomical (aCompCor). For aCompCor, three probabilistic masks (CSF, WM and combined CSF+WM) are generated in anatomical space. The implementation differs from that of (Behzadi et al., 2007) in that instead of eroding the masks by 2 pixels on BOLD space, the aCompCor masks are subtracted a mask of pixels that likely contain a volume fraction of GM. This mask is obtained by dilating a GM mask extracted from the FreeSurfer’s *aseg* segmentation, and it ensures components are not extracted from voxels containing a minimal fraction of GM. Finally, these masks are resampled into BOLD space and binarized by thresholding at 0.99 (as in the original implementation). Components are also calculated separately within the WM and CSF masks. For each CompCor decomposition, the *k* components with the largest singular values are retained, such that the retained components’ time series are sufficient to explain 50 percent of variance across the nuisance mask (CSF, WM, combined, or temporal). The remaining components are dropped from consideration. The head-motion estimates calculated in the correction step were also placed within the corresponding confounds file. The confound time series derived from head motion estimates and global signals were expanded with the inclusion of temporal derivatives and quadratic terms for each (Satterthwaite et al., 2013). Frames that exceeded a threshold of 0.5 mm FD or 1.5 standardized DVARS were annotated as motion outliers. The BOLD time-series were resampled into standard space, generating a *preprocessed BOLD run in MNI152NLin2009cAsym space*. Many internal operations of *fMRIPrep* use *Nilearn* 0.8.1 ((Abraham et al., 2014), RRID:SCR_001362), mostly within the functional processing workflow. For more details of the pipeline, see <u>the section corresponding to workflows in *fMRIPrep*’s documentation</u>.

For the streamlined application of additional noise components and data-cleaning strategies within a single framework, we utilized rs-Denoise ((Dubois et al., 2018), see https://github.com/adolphslab/rsDenoise), an open-source Python-based pipeline. This pipeline involved several steps: (1) z-score normalization of the signal at each voxel; (2) removal of linear and quadratic trends with polynomial regressors; (3) utilization of *fMRIPrep’s aCompCor* parameters, to regress out five components derived from whole-brain mean signals; (4) utilization of translational and rotational realignment parameters and their temporal derivatives as explanatory variables in motion regression; (5) temporal filtering was performed with a discrete cosine transform (DCT) filter with a cutoff frequency of 0.008 Hz. Lastly, the pre-processed runs were smoothed using a 4-mm full-width at half maximum (FWHM) Gaussian kernel and merged on the temporal domain.

### Data analysis

#### Amygdala-prefrontal cortex functional connectivity

First, binary masks of the two regions of interest were generated. For this purpose, a parcellation atlas that combined cortical (400 Parcels and 7 Networks) and subcortical (Scale 1) parcellations was employed to delineate the amygdala and prefrontal cortex masks (Tian et al., 2020). Following this, since the data were already cleaned by regressing out the confounds of interest, the BOLD eigenvariate specific to the amygdala region was extracted as region average signal using the "fslmeants" command. Afterward, the extracted amygdala average signal was used as a regressor in a general linear model to correlate, per subject, the average amygdala activity with each prefrontal cortex voxel. Spatial maps for every subject were generated from the last step and merged into a 4D volume that was subsequently used as input to FSLrandomise (FSL version 6.0.3). Randomization, with ten thousand permutations, was used to associate the nightmare frequency scores to its participant functional connectivity map. The GLM included nightmare frequency as the main regressor of interest, as well as weekly dream frequency scores, sex and age as confound regressors.. After permutations, *FSL randomise* outputs a Threshold-Free Cluster Enhancement (TFCE) map corrected for multiple comparisons. TFCE aims to preserve the sensitivity advantages of cluster-based inference while avoiding arbitrary cluster-forming threshold. This approach yields an output image at the voxel level, where each voxel’s value represents the accumulative cluster-like local spatial support at a range of cluster-forming thresholds (Salimi-Khorshidi et al., 2011; Smith & Nichols, 2009).

#### ReHo analysis

Single ReHo maps were generated by calculating Kendall’s coefficient of concordance (KCC). This metric assesses the regional homogeneity of the blood oxygen level dependent time series within each voxel and its 26 adjacent voxels. The generation process used the *3dReHo* function in *AFNI* (versions 22.1.09 and 23.0.02 for Datasets 1 and 2, respectively) (Zang et al., 2004). Subsequently, the individual ReHo maps were normalized by dividing the KCC in each voxel by the mean KCC of the whole gray matter. Finally, the ReHo maps were smoothed using a 4-mm full-width at half maximum (FWHM) Gaussian kernel.

We adopted a dual approach to the statistical analysis. First, we examined differences between two groups: high nightmare sufferers (at least one nightmare per week) and matched controls (less than one per year), as a direct replication of the previous literature, by pooling the two extreme groups from the combination of Dataset 1+2. Second, we examined parametric differences across the spectrum of nightmare frequency in a large dataset derived by combining Dataset 1+2 (see Table 1). For both the group-comparison replication analysis and continuous nightmare frequency scores, we used a more stringent threshold of *p*<0.001 at the voxel level, contrary to the significance threshold from Marquis et al. and Shen et al. (Marquis et al., 2021; Shen et al., 2016), set at *p*<0.01 at the voxel level. It is important to note that choosing a critical statistical threshold (CDT) of 0.01 has been shown to yield excessive false positives (see (Eklund et al., 2016) for a detailed discussion). Cluster-level threshold values were estimated in *SPM*. Because parametric statistical methods for group analysis, such as *SPM*, can produce erroneously low FWE-corrected cluster p-values, thereby inflating statistical significance, we also used *FSL randomise* as a nonparametric method to evaluate our results (Eklund et al., 2016). Dream recall frequency, age, sex, and site were controlled for in the ReHo analyses. Statistical analyses were performed using *SPM12* (Statistical Parametric Mapping 12, Wellcome Trust Centre for Neuroimaging, Institute of Neurology, University College London, United Kingdom) with *Matlab* (R2022a, The Mathworks, Natick, MA, United States).

## Results

### Demographics and questionnaires

Participants in Dataset 1 reported an average dream recall frequency of 2.18 times per week (*SD*=2.05) and experienced nightmares 1.34 times per month (*SD*=3.31), with a frequency ranging from once to several times per week. For Dataset 2, participants reported an average dream recall frequency of 2.10 times per week (*SD*=2.16) and nightmares at an average frequency of 1.02 times per month (*SD*=2.83). In partial agreement with a higher incidence of nightmares in females suggested in previous literature (Nielsen & Levin, 2007), for Dataset 1 the data indicated significant differences in nightmare frequency between females (n=152, mean=1.61, *SD*=3.51) and males (n=108, mean=0.96, *SD*=2.97), *W*=5782, *p*<0.001 (two-tailed), but not in dataset 2 (n=99 females: mean=1.28, *SD*=3.56; vs. n=65 males: mean =0.67, SD=0.99), *W* =2939, *p*=0.33 (two-tailed). There was no evidence of age-related differences in dream recall frequency (Dataset 1: rho=-0.052, *p*=0.40; Dataset 2: rho=-0.026, *p*=0.74) or nightmare frequency (Dataset 1: rho=-0.059, *p*=0.347; Dataset 2: rho=-0.037, *p*=0.638). As expected, dream recall and nightmare frequency were significantly correlated in both Dataset 1 (rho=0.338, *p*<0.001) and Dataset 2 (rho=0.216, *p*<0.005) (Figure 1). Nevertheless, sex, age, and dream recall frequency were controlled for in all analyses. When comparing the HNF group to the CTL group, a higher dream recall frequency was observed in the HNF group but there were no significant age or sex differences between the two groups (*p*>0.66).

**Figure 1:**
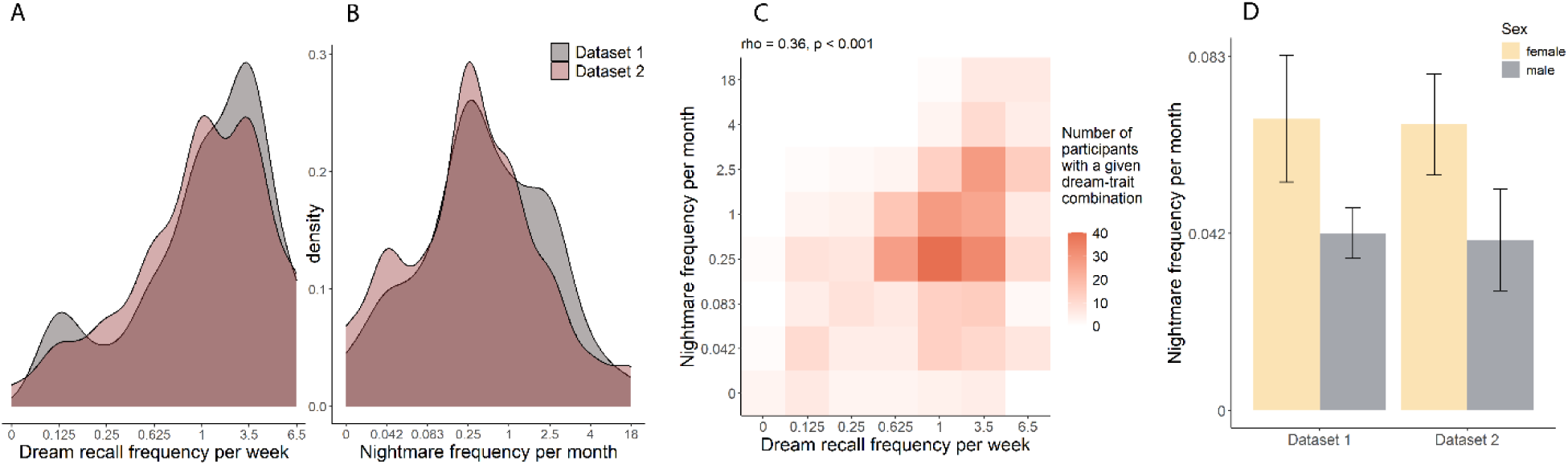
Overview of the behavioral data in the two Datasets. **A:** Density distributions of the dream recall frequency scores (recoded into units per week) for Dataset 1 and Dataset 2; **B:** Density distributions of the nightmare frequency scores (recoded into units per month) for Dataset 1 and Dataset 2; **C:** A heatmap for the combination of nightmare frequency and dream recall frequency scores across the two Datasets, and the correlation between the scores. Darker colors denote more frequent combinations; **D:** Average nightmare frequency scores for male and female participants, for both Dataset 1 and Dataset 2. The vertical bars denote standard errors.

### Amygdala-prefrontal cortex functional connectivity relationship with nightmare frequency

In our investigation of functional connectivity between the amygdala and prefrontal cortex and its relationship to nightmare frequency, we first analyzed Dataset 1. This initial analysis revealed no significant voxels (*p_FWEc_* = 0.67). To validate these findings, we replicated the analysis using an independent Dataset 2, which also showed no significant results (*p_FWEc_* = 0.65). In other words, no functional connectivity between these regions were statistically significantly associated with nightmare frequency within the parameters of our study. Similar results were found when all analyses were repeated using raw nightmare frequency scores (Supplementary Figure 1).

### ReHo analysis

We performed a ReHo analysis to explore potential group differences between high and low nightmare frequency, as previously reported in the literature. The results showed no significant differences in ReHo scores between groups (Supplementary Tables 1 and 2). However, when examining continuous nightmare frequency scores across the combined Dataset 1+2, we identified a significant cluster in the cerebellum (peak-voxel t-value=5.87, MNI coor=24,-68,-60, Figure 2a).

**Figure 2:**
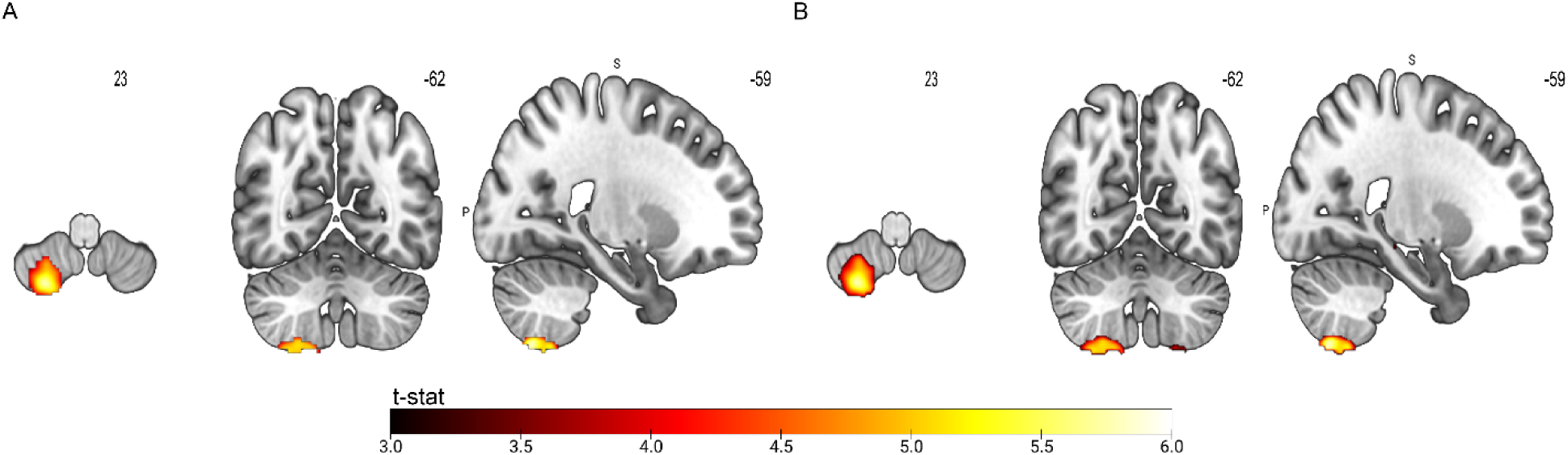
Regional homogeneity analyses results: **A:** Dataset 1+2 combined (N=424) SPM parametric analysis resulted in a significant cluster located in the cerebellum (peak-voxel t-value=5.87, MNI coor=24,-68,-60); **B:** Dataset 1+2 combined (N=424) non-parametric FSL Randomise resulted in a similar significant cluster located in the cerebellum (peak-voxel t-value=5.96, MNI coor=23.9,-67.5,-59.7).

To robustly test these findings, we used a nonparametric permutation test in addition to a threshold-free cluster enhancement approach. This rigorous analysis revealed no significant clusters in the group comparison (Supplementary Figure 2). However, in the analysis of the continuous nightmare frequency score, we identified a single significant cluster in the cerebellum (cluster size=150 voxels; *p_FWEc_*<0.01; MNI coor=23.9,-67.5,-59.7) This finding is shown in Figure 2b. In addition, we repeated all analyses using raw nightmare frequency scores (i.e. the eight-point rating scale instead of the recoded monthly scale), which did not yield significant results (for group comparison, see Supplementary Tables 3 and 4, Supplementary Figure 3; for continuous nightmare frequency scores see: Supplementary Tables 5, Supplementary Figure 4).

## Discussion

The present study had two main aims. First, to investigate the relationship between nightmare frequency and functional connectivity between two key regions, amygdala and prefrontal cortex, involved in emotional regulation and fear extinction processes, and second, to replicate the existing literature on the neural correlates of nightmares in two large study cohorts. Contrary to our initial hypothesis, our analysis did not reveal a significant relationship between nightmare frequency and functional resting connectivity between the prefrontal cortex and the amygdala. In addition, while we were able to partially replicate previous whole-brain ReHo findings on nightmare frequency, these findings did not withstand rigorous tests with more appropriate statistical approaches. Despite our increased statistical power compared to previous studies, we could reliably identify only a single cluster located in the cerebellum and only when analyzing nightmare frequency scores continuously (as opposed to grouped scores as in previous studies).

Building on the sensory processing sensitivity framework for nightmares (Carr & Nielsen, 2017), we hypothesized that connectivity between the amygdala and prefrontal cortex might serve as a potential predictor of nightmare frequency. This hypothesis stemmed from previous studies linking such coupling to psychiatric conditions and disorders such as anxiety levels (Kim et al., 2011), anxiety disorders (Prater et al., 2013), major depressive disorder (Tang et al., 2013), post-traumatic stress disorder (Sripada et al., 2012), dream emotionality (Blake et al., 2019), and threat perception (Ochsner & Gross, 2005). Our rationale was based on the expectation that heightened emotional reactivity will manifest in resting-state amygdala-prefrontal connectivity, especially given the known association between prefrontal cortex activation and nightmare severity/distress during negative emotional stimulation (Carr, 2020; Marquis et al., 2019; Sterpenich et al., 2020). Contrary to our predictions, our analysis of Dataset 1 and its independent replication in Dataset 2 did not yield statistically significant results. Although we did not expect nightmare formation to depend solely on amygdala-prefrontal cortex connectivity, we considered this to be the most prominent candidate for a trait correlate of nightmare frequency, considering the crucial role these regions play in relevant processes and existing models of nightmare formation. However, the lack of a significant relationship in our study despite a large sample size in two datasets may suggest that any potential association might be subtle if existent, potentially overshadowed by individual differences confounders such as personality traits including sensory processing sensitivity, or trauma history.

In investigating ReHo in the neural correlates of nightmares, we attempted to replicate previous group comparison methodology while addressing concerns related to the reproducibility of fMRI studies. Shen et al. (Shen et al., 2016) found elevated ReHo in the left anterior cingulate cortex and right inferior parietal lobule in patients with nightmare disorder, while Marquis et al. (Marquis et al., 2021) found altered ReHo in several brain regions. In contrast to Shen et al., Marquis et al. did not find group differences in ReHo in the anterior cingulate cortex or inferior parietal lobule, and the studies had different sample populations. Shen et al. focused on patients with nightmare disorder and a strict group of healthy controls with minimal lifetime nightmares. In contrast, Marquis et al. included a mix of high nightmare sufferers and bad dream recallers with no awakenings after disturbing dreams. Population differences may have contributed to the contrasting results. Our study aimed to replicate the group comparison by focusing on high nightmare sufferers and realistic healthy controls. We calculated the group comparison analysis in two ways to control false positive rates: 1) using a similar approach and CDT (*p*<0.001) but using *SPM* to estimate the cluster extension value, and 2) given the inflated statistical inference of parametric methods, we used a nonparametric permutation approach as implemented in *FSL Randomise*. No significant clusters survived these additional approaches. The fMRI community has faced a reproducibility problem in part because low CDT can produce misleadingly low cluster P-values, raising concerns about the accuracy of many published fMRI studies (Eklund et al., 2016). Nevertheless, previous work has used a CDT=0.01, which is known to result in higher false-positive rates. We performed analyses analogous to those used for group comparisons to examine the neural correlates of nightmares across participants’ continuous severity levels. Using a parametric (alpha-level voxel thresholding followed by Gaussian random field theory) and non-parametric (TFCE followed by permutation testing) cluster inference approaches, only the cerebellum cluster survived corrections. Previous studies have reported decreased ReHo values in the cerebellum in individuals with nightmare experiences, which is consistent with our findings (Marquis et al., 2021).

The cerebellum, traditionally associated with motor functions and considered less functionally significant than the cerebral cortex, in recent years has attracted increased attention also with respect to cognitive and emotional processing (Adamaszek et al., 2022; Baillieux et al., 2008; Sacchetti et al., 2009). For example, the cerebellum has been implicated in the formation, consolidation, and extinction of fear memories and other emotion modulations (Rudolph et al., 2023). Although poorly characterized in sleep and largely unexplored in nightmares, the cerebellum is linked to anxiety disorders (Moreno-Rius, 2018), major depression disorder (Fitzgerald et al., 2008), and bipolar disorder (Hoppenbrouwers et al., 2008), which are all associated with nightmare distress and frequency. Anatomically, cerebellar connections to the limbic system suggest its potential role in the brain’s emotional network (Çavdar et al., 2018; Hilber et al., 2019; Novello et al., 2024), and a recent study has demonstrated direct monosynaptic projections from the cerebellum to the amygdala (Zhang et al., 2024). Further cerebellar projections target the thalamus and the brainstem, which are implicated in REM sleep regulation (Sathyanesan et al., 2019), and also parts of the cerebellum have been shown to be activated during REM sleep (Braun, 1997; Canto et al., 2017; Sokoloff et al., 2015). Of note, anxiolytic benefits of physical activity have been correlated with increased activity of the cerebellar projections to the amygdala (Zhang et al., 2024), which is highly activated during REM sleep (Corsi-Cabrera et al., 2016; Maquet, 1997; Nofzinger et al., 1997). Considering that REM sleep dream narratives are characterized by a high level of experienced motor activity (Porte & Hobson, 1996), it is tempting to speculate that cerebellum-amygdala projections play a role in emotionally arousing dream content such as nightmares. To test this possibility, we added an analysis to probe the association between nightmare frequency and cerebellar-amygdala functional connectivity, using the results of the ReHo analysis to define a relevant cerebellar region of interest (See Supplementary Material for details on methods). No significant functional relationship between the amygdala and our specific cerebellar region was find, yet the above-mentioned results indicate that the cerebellum remains an important structure for future exploration, considering also the recent discovery of an amygdala-independent pathway for fear processing (Wang et al., 2024). Accordingly, the precise involvement of the cerebellum and interactions with other brain regions in the domains of emotion regulation processes and dream emotionality warrants further investigation.

Several limitations of our study have to be considered. First, the questionnaire assessing dreaming and nightmare frequency may have influenced the present results as they did not allow to check for levels of nightmare distress. Moreover, as our cohorts consisted of healthy young participants, generalizations to patients or the entire population are difficult. In addition, we carefully considered the methodological parameters for the ReHo analysis, guided by previous research (Maximo et al., 2013) and the literature we sought to replicate. Spatial smoothing was performed after ReHo computations to prevent inflation of correlation statistics by averaging signals over a larger area. Because ReHo measures local connectivity, the choice of neighborhood size for ReHo analysis is another important consideration, allowing researchers to specify the extent of the neighborhood to be tested for correlation with each voxel. This study chose a neighborhood size of 27 voxels to replicate findings from previous literature (Marquis et al., 2021; Shen et al., 2016). In particular, choosing a neighborhood size of 27 voxels means that the analysis considers every voxel with an adjacent face, edge, or corner. This may explain why the significant cluster accounted for the edge of the cerebellum despite its proximity to the brain boundary. In spite of these shortcomings, it is clear that the large sample size and robust threshold correction methods are strong points to support the outcomes of our study.

The combination of dream research and neuroimaging comes with several challenges, as it is impossible to time-lock the exact time a dream experience occurred, and nightmares are rare in sleep laboratories and may imply major imaging motion artifacts. Future research should address these design limitations by combining dream diaries and retrospective questionnaires with all-night imaging recordings. A dream diary is an established method for evaluating dream content, however, it may overestimate dreaming and nightmare frequency as a continued dream diary enhances dream recollection (Stumbrys et al., 2013). In contrast, retrospective questionnaires have been shown to underestimate dreaming and nightmare frequency (Wood & Bootzin, 1990). Accordingly, a combination of the two approaches might give a more robust assessment of the occurrence of nightmares in the study population.

## Conclusion

In summary, contrary to our initial expectations, we did not find a significant relationship between nightmare frequency and functional connectivity between the prefrontal cortex and the amygdala, key regions involved in emotional regulation and fear extinction processes. In contrast, probing the relationship between nightmare frequency and regional homogeneity in a whole-brain analysis, we did find a role of the cerebellum in nightmare frequency, supporting an increasingly discussed role of the cerebellum in emotional processing. Functional connectivity of this cerebellar region with the amygdala, however, was not associated with nightmare frequency.

While our study replicated the group comparison methodology used by Shen et al. and Marquis et al., the complexities surrounding the reproducibility of fMRI studies should be considered. Our efforts to control false-positive rates through various recalculations, including stringent cluster-defining thresholds and nonparametric permutation approaches, did not yield significant clusters except in the cerebellum, highlighting the intricacies involved in interpreting neuroimaging data in general and our specific findings. The unexpected lack of robust significant results, especially given the larger sample size in the current study, prompts a reevaluation of existing models and emphasizes the need to account for individual differences, such as personality traits, trauma history, and cognitive processes. As we navigate the complexity of neural circuits and brain regions involved in nightmares, these findings contribute to the ongoing dialogue in the field, fostering a deeper understanding of the neurobiology behind nightmares and guiding future research efforts.

## Supplementary Material

Supplementary material is available in a separate file.

## Acknowledgments

This article is based upon work fromCOST Action CA18106, The Neural Architecture of Consciousness, supported by COST (The European Cooperation in Science and Technology). MP and MD were supported by a Vidi grant from the Dutch Research Council (NWO) and a research grant from the Bial Foundation. We thank Dunja Paunovic, Katarina Vulic, Blanka Zana, Paola Galdi, Katarzyna Hat for their enormous efforts during data collection, quality checks, and pre-processing. We also thank Nils Muller, who helped contextualize the analysis.

## Data availability

Data will be available upon reasonable request.

