## Supplementary Figure 1 for "Neural correlates of nightmares revisited: findings from large-scale fMRI cohorts"

**Functional connectivity relationship between amygdala-prefrontal cortex and nightmare frequency**

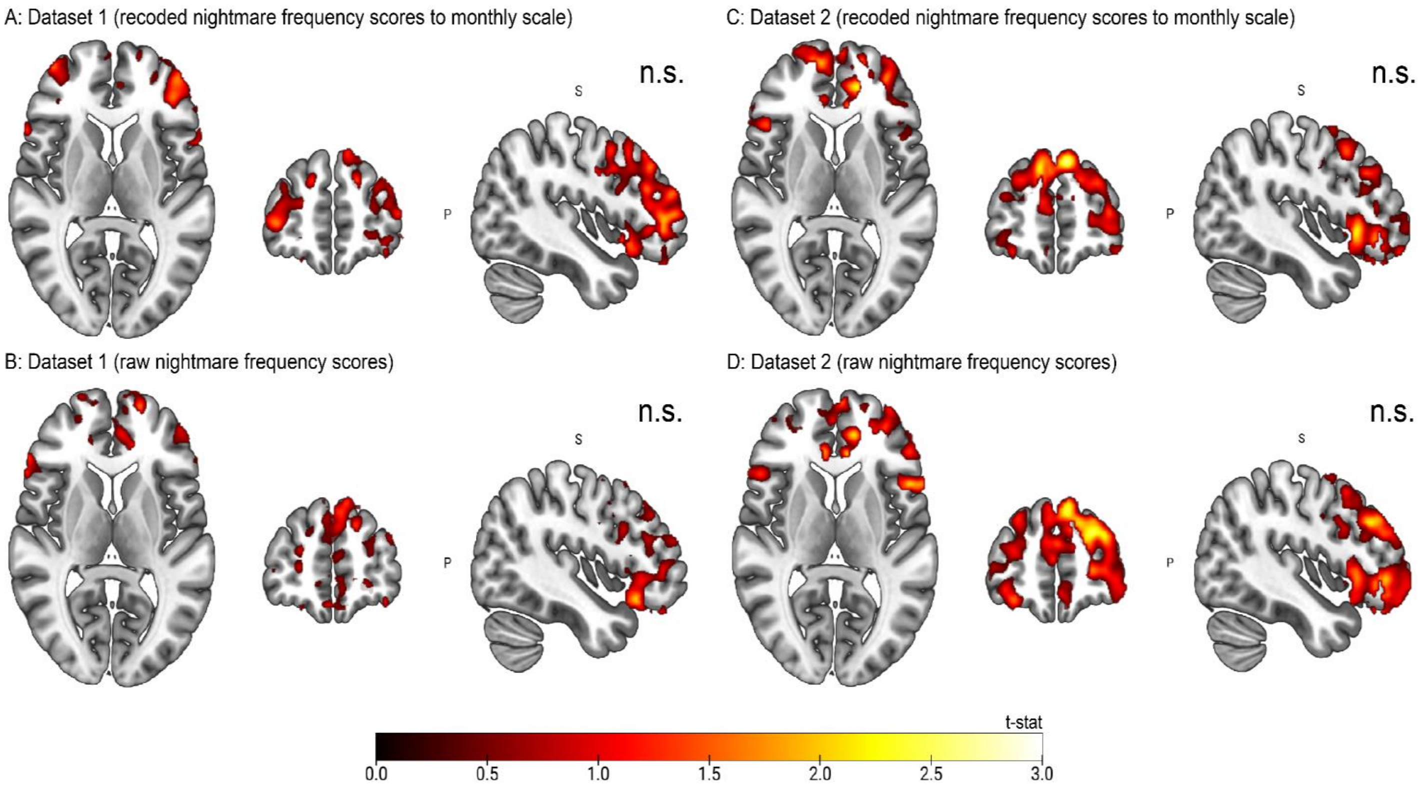


**Supplementary Figure 1:** Amygdala-prefrontal cortex functional connectivity: Dataset 1 (N=260) resulted in non-significant clusters a) by using the recoded nightmare frequency scores to a monthly scale (Stumbrys et al., 2013) ($p_{FWEc}$ = 0.67), and b) by using the raw nightmare frequency scores ($p_{FWEc}$ = 0.59). Dataset 2 (N=164) also resulted in non-significant clusters c) by using the recoded nightmare frequency scores to a monthly scale ($p_{FWEc}$ = 0.48), and d) by using the raw nightmare frequency scores ($p_{FWEc}$ = 0.36). All t-maps are in MNI coor=-44,53,7). Please note that the here shown activations are of descriptive nature only (thresholded at $p<0.05$) and do not reflect significant results.
