## Supplementary Table 1 for "Neural correlates of nightmares revisited: findings from large-scale fMRI cohorts"

**Regional homogeneity (ReHo) analysis**

**A: Group comparison replication analyses with recoded nightmare frequency scores on a monthly scale:**

- Parametric results (*SPM*):

**Supplementary Table 1:** High nightmare frequency > healthy controls: p-values adjusted for search volume.

| cluster-level | | | peak-level | | | |
|---|---|---|---|---|---|---|
| $p_{FWEc}$ | $q_{FDRc}$ | $K_E$ | $T$ | x (mm) | y (mm) | z (mm) |
| 0.999 | 0.859 | 8 | 4.39 | 66 | -22 | 44 |
| 0.999 | 0.859 | 6 | 3.85 | -38 | -40 | -50 |
| 0.995 | 0.859 | 13 | 3.68 | -48 | -44 | 42 |
| 0.999 | 0.859 | 7 | 3.67 | 38 | -58 | -62 |
| 1.000 | 0.859 | 5 | 3.62 | -52 | 44 | -14 |
| 1.000 | 0.859 | 4 | 3.58 | 50 | -58 | -52 |
| 1.000 | 0.859 | 4 | 3.55 | 10 | -38 | 34 |
| 1.000 | 0.859 | 2 | 3.47 | -26 | -16 | -36 |
| 1.000 | 0.859 | 4 | 3.44 | 54 | -68 | -18 |
| 1.000 | 0.859 | 1 | 3.34 | 10 | -52 | 54 |
| 1.000 | 0.859 | 1 | 3.33 | 48 | -50 | -52 |
| 1.000 | 0.859 | 1 | 3.32 | -42 | -50 | -58 |
| 1.000 | 0.859 | 1 | 3.32 | 34 | -74 | -58 |
