## Supplementary Table 2 for "Neural correlates of nightmares revisited: findings from large-scale fMRI cohorts"

**Regional homogeneity (ReHo) analysis**

**A: Group comparison replication analyses with recoded nightmare frequency scores on a monthly scale:**

- Parametric results (*SPM*):

**Supplementary Table 2:** High nightmare frequency < healthy controls: p-values adjusted for search volume.

| cluster-level | | | peak-level | | | |
|---|---|---|---|---|---|---|
| $p_{FWEc}$ | $q_{FDRc}$ | $K_E$ | $T$ | x (mm) | y (mm) | z (mm) |
| 0.467 | 0.339 | 86 | 6.17 | -58 | -22 | 10 |
| 0.284 | 0.339 | 115 | 4.40 | -60 | -30 | -18 |
| 0.830 | 0.586 | 45 | 4.30 | 4 | -88 | 18 |
| 0.981 | 0.709 | 20 | 4.21 | -14 | -40 | 78 |
| 0.886 | 0.586 | 38 | 3.75 | 14 | 44 | 10 |
| 0.999 | 0.834 | 7 | 3.71 | -4 | 44 | 16 |
| 1.000 | 0.859 | 2 | 3.70 | -44 | 8 | 58 |
| 0.971 | 0.709 | 23 | 3.63 | -50 | -44 | -22 |
| 0.999 | 0.834 | 7 | 3.57 | 10 | 24 | 34 |
| 0.999 | 0.834 | 6 | 3.50 | -18 | -8 | -18 |
| 1.000 | 0.834 | 5 | 3.49 | -4 | -60 | 0 |
| 1.000 | 0.859 | 1 | 3.34 | -54 | -16 | 54 |
| 1.000 | 0.859 | 1 | 3.32 | 4 | 56 | 12 |
