## Supplementary Figure 2 for "Neural correlates of nightmares revisited: findings from large-scale fMRI cohorts"

**Regional homogeneity (ReHo) analysis**

**A: Group comparison replication analyses with recoded nightmare frequency scores on a monthly scale:**

- Non-parametric results (*FSL randomise*):

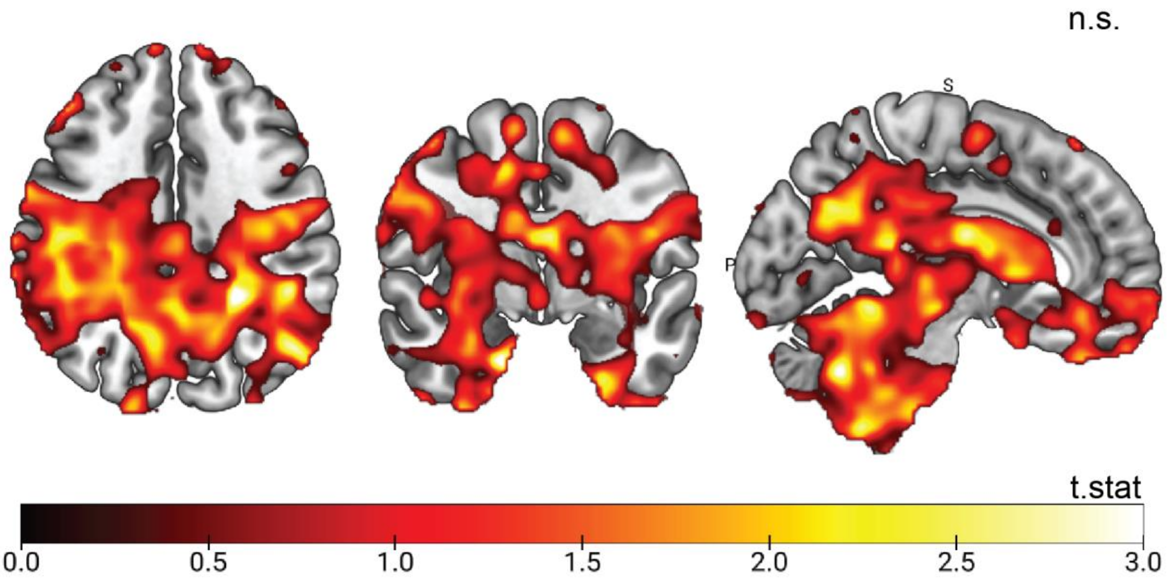


**Supplementary Figure 2:** Group comparison replication analysis, high vs. low nightmare frequency score using the recoded nightmare frequency scores to a monthly scale ($p_{FWEc}$ = 0.294) (Stumbrys et al., 2013). The t-map is in MNI coor=-6,-3.8,37). Please note that the here shown activations are of descriptive nature only (thresholded at $p<0.05$) and do not reflect significant results.
