## Supplementary Table 3 for "Neural correlates of nightmares revisited: findings from large-scale fMRI cohorts"

**Regional homogeneity (ReHo) analysis**

**B: Group comparison replication analyses with raw nightmare frequency scores:**

- Parametric results (*SPM*):

**Supplementary Table 3:** High nightmare frequency > healthy controls: p-values adjusted for search volume

| cluster-level | | | peak-level | | | |
|---|---|---|---|---|---|---|
| $p_{FWEc}$ | $q_{FDRc}$ | $K_E$ | $T$ | x (mm) | y (mm) | z (mm) |
| 1.000 | 0.860 | 4 | 3.87 | 66 | -22 | 44 |
| 0.990 | 0.860 | 16 | 3.81 | 38 | -58 | -62 |
| 0.998 | 0.860 | 9 | 3.53 | 2 | 44 | -18 |
| 1.000 | 0.860 | 3 | 3.46 | 52 | -68 | -18 |
| 1.000 | 0.860 | 2 | 3.43 | 46 | 14 | -42 |
| 1.000 | 0.860 | 5 | 3.41 | 22 | -40 | -52 |
| 1.000 | 0.860 | 2 | 3.36 | -40 | -42 | -50 |
| 1.000 | 0.860 | 1 | 3.32 | 48 | 46 | -18 |
