## Supplementary Table 4 for "Neural correlates of nightmares revisited: findings from large-scale fMRI cohorts"

**Regional homogeneity (ReHo) analysis**

**B: Group comparison replication analyses with raw nightmare frequency scores:**

- Parametric results (*SPM*):

**Supplementary Table 4:** High nightmare frequency < healthy controls: p-values adjusted for search volume

| cluster-level | | | peak-level | | | |
|---|---|---|---|---|---|---|
| $p_{FWEc}$ | $q_{FDRc}$ | $K_E$ | $T$ | x (mm) | y (mm) | z (mm) |
| 0.346 | 0.195 | 104 | 7.13 | -58 | -22 | 10 |
| 0.166 | 0.167 | 146 | 4.31 | -60 | -30 | -16 |
| 0.939 | 0.843 | 30 | 4.11 | 12 | 24 | 34 |
| 0.990 | 0.843 | 16 | 3.90 | -18 | -36 | 76 |
| 0.983 | 0.843 | 19 | 3.90 | 4 | -90 | 18 |
| 1.000 | 0.860 | 3 | 3.49 | -56 | -18 | 52 |
| 1.000 | 0.860 | 5 | 3.49 | -44 | -52 | -46 |
| 1.000 | 0.860 | 5 | 3.47 | -50 | -40 | 24 |
| 1.000 | 0.860 | 3 | 3.43 | -30 | -60 | 58 |
| 1.000 | 0.860 | 2 | 3.40 | -46 | -16 | 48 |
| 1.000 | 0.860 | 1 | 3.32 | -48 | -18 | 50 |
