## Supplementary Figure 3 for "Neural correlates of nightmares revisited: findings from large-scale fMRI cohorts"

**Regional homogeneity (ReHo) analysis**

**B: Group comparison replication analyses with raw nightmare frequency scores:**

- Non-parametric results (*FSL randomise*):

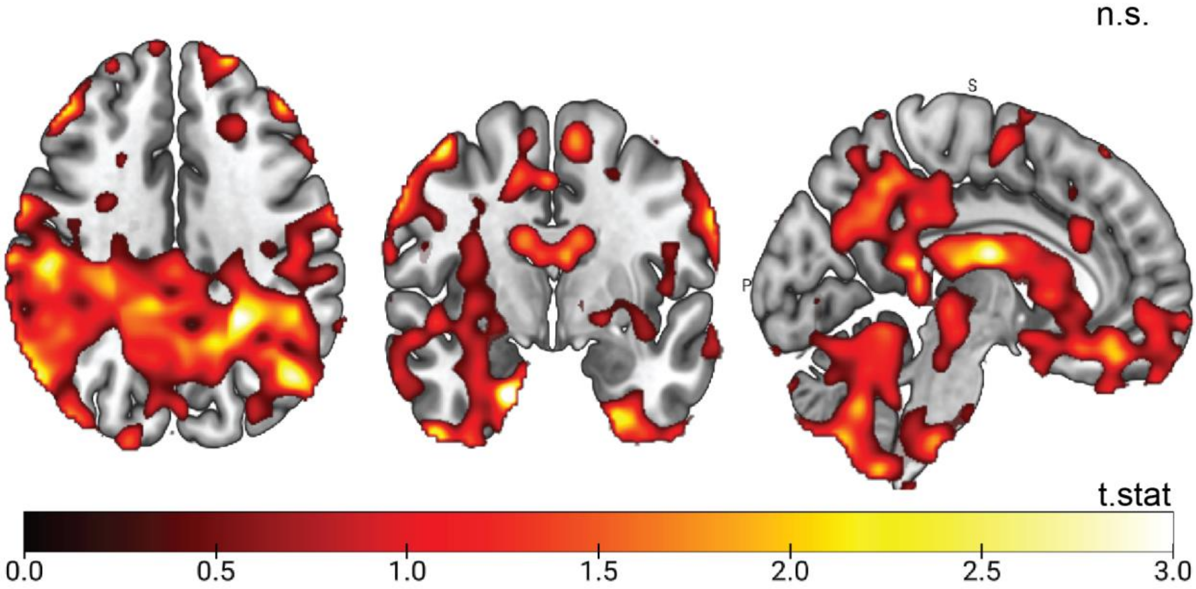


**Supplementary Figure 3:** Group comparison replication analysis, high vs. low nightmare frequency score using the raw nightmare frequency scores ($p_{FWEc}$ = 0.844). The t-map is in MNI coor=-6,-3.8,37). Please note that the here shown activations are of descriptive nature only (thresholded at $p<0.05$) and do not reflect significant results.
