## Supplementary Table 5 for "Neural correlates of nightmares revisited: findings from large-scale fMRI cohorts"

### Regional homogeneity (ReHo) analysis

#### C: Parametric results of combined Datasets 1+2 (N=464) using raw nightmare frequency scores:

- Parametric results (*SPM*):

**Supplementary Table 5:** Statistics: p-values adjusted for search volume.

| cluster-level | | | peak-level | | | |
|---|---|---|---|---|---|---|
| $p_{FWEc}$ | $q_{FDRc}$ | $K_E$ | $T$ | x (mm) | y (mm) | z (mm) |
| 0.977 | 0.875 | 20 | 4.02 | -50 | 46 | 12 |
| 0.948 | 0.875 | 28 | 3.53 | -40 | -14 | 24 |
| 0.997 | 0.875 | 9 | 3.31 | -46 | -6 | 6 |
| 1.000 | 0.875 | 2 | 3.27 | 62 | -32 | 50 |
| 0.999 | 0.875 | 5 | 3.27 | 38 | -22 | -36 |
| 1.000 | 0.875 | 1 | 3.25 | 46 | -52 | 58 |
| 0.999 | 0.875 | 4 | 3.25 | -56 | 38 | 2 |
| 1.000 | 0.875 | 1 | 3.13 | 44 | 16 | -34 |
| 0.999 | 0.875 | 5 | 3.13 | -58 | 24 | 16 |
| 1.000 | 0.875 | 2 | 3.13 | 34 | -62 | -62 |
| 1.000 | 0.875 | 1 | 3.11 | -40 | 40 | 2 |
