## Supplementary Figure 4 for "Neural correlates of nightmares revisited: findings from large-scale fMRI cohorts"

**Regional homogeneity (ReHo) analysis**

**C: Parametric results of combined Datasets 1+2 (N=464) using raw nightmare frequency scores:**

- Non-parametric results (*FSL randomise*):

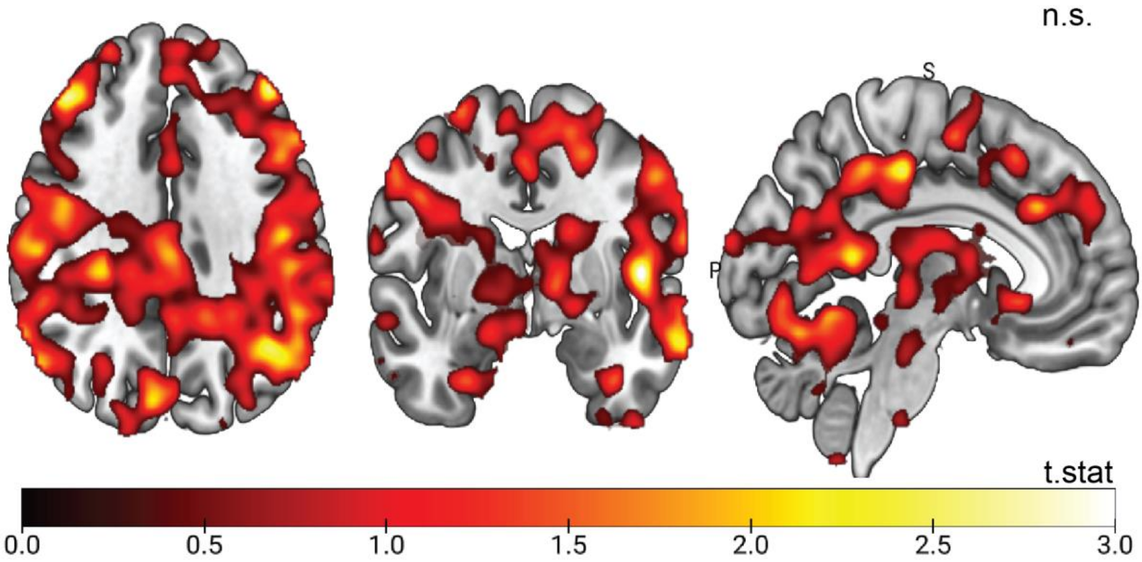


**Supplementary Figure 4:** ReHo analysis combining Datasets 1+2 using the raw nightmare frequency scores ($p_{FWEc}$ = 0.676). The t-map is in MNI coor=-6,-3.8,37). Please note that the here shown activations are of descriptive nature only (thresholded at p<0.05) and do not reflect significant results.
